# Performance Determinants of Unsupervised Clustering Methods for Microbiome Data

**DOI:** 10.1101/2021.04.08.439060

**Authors:** Yushu Shi, Liangliang Zhang, Christine Peterson, Kim-Anh Do, Robert Jenq

## Abstract

**Background:** In microbiome data analysis, unsupervised clustering is often used to identify naturally occurring clusters, which can then be assessed for associations with characteristics of interest. In this work, we systematically compared beta diversity and clustering methods commonly used in microbiome analyses. We applied these to four published datasets where highly distinct microbiome profiles could be seen between sample groups.

**Results:** Although no single method outperformed the others consistently, we did identify key scenarios where certain methods can underperform. Specifically, the Bray Curtis metric resulted in poor clustering in a dataset where high-abundance OTUs were relatively rare. In contrast, the unweighted UniFrac metric clustered poorly when used on a dataset with a high prevalence of low-abundance OTUs. To test our proposition, we systematically modified properties of the poorly performing datasets and found that this approach resulted in improved Bray Curtis and unweighted UniFrac performance. Based on these observations, we rationally combined the Bray Curtis metric and the unweighted UniFrac metrics and found that this new beta diversity metric showed high performance across all datasets. We also evaluated our findings by examining a clinical dataset where clusters are less separated.

**Conclusions:** Our systematic evaluation of clustering performance in these five datasets demonstrates that there is no existing clustering method that universally performs best across all datasets. We propose a combined metric of Bray Curtis and unweighted UniFrac that capitalizes on the complementary strengths of the two metrics.

## Background

The development of next generation sequencing in the past decade has increased access for researchers to analyze microbial communities [8, 13]. A common method involves deep sequencing of 16S rRNA genes and grouping bacteria at a certain level of 16S rRNA gene similarity [11]. The highest resolution bacterial sequence is referred to as an operational taxonomic unit (OTU). A challenge with microbiome analyses is that these sequences are derived from bacteria that can be incredibly diverse, and simultaneously have a relatedness structure that is internally associated, both genetically and phenotypically. Metrics to account for phylogenetic closeness have been proposed specifically for microbiome data [15, 16, 3]. Previously, Fukuyama [7] showed that deep and shallow parts of the tree contribute differently to phylogenetic distances.

Microbiome data analysis can suffer from overgeneralization (e.g., comparing observations by alpha diversity measures such as Shannon [22] or Simpson [24] diversity) or overspecification (e.g., performing association tests for each taxon, which incurs a heavy penalty for multiple testing). A useful and common strategy to address both of these limitations is to first apply an unsupervised clustering methodology. After considering the microbiome data as a whole and identifying naturally occurring clusters of samples, these clusters can then be assessed for associations with sample characteristics of interest. Previous studies, for instance, have shown that human stool microbiome samples naturally form clusters that are associated with dietary and geographic factors [21, 17]. A problem that often confuses researchers is that clustering performance results often vary depending on the algorithm or the beta diversity metrics used, observed previously by Koren, et al. [14] and Claesson, et al. [4].

When performing microbiome sample clustering, both model-based methods and machine learning methods have been used. Machine learning methods, which rely on defined distance metrics, are used more frequently than model-based statistical methods, due to their efficient implementation and easy interpretation. In this paper, we focused on the Partition Around Medoids (PAM) [12] clustering method, which is related to but considered more robust than K-means. In contrast to K-means, which can be sensitive to the effects of outliers, PAM’s optimization goal is to minimize the sum of distances to the medoids instead of minimizing the sum of the squared distances to the cluster centers. We also evaluated another frequently used method for non-Euclidean metrics, hierarchical clustering with complete linkage. This method initially bundles the closest observations (with distances defined by the longest distance between any two observations in two clusters), and gradually results in a binary tree that combines the two closest clusters.

Supplemental Figure S1 illustrates differences between these clustering methods. The choice of clustering algorithm and distance metric together determine the performance of a machine learning clustering method. The metrics considered in this study are commonly used and include the Bray-Curtis dissimilarity metric [2], the unweighted UniFrac distance [15], the weighted UniFrac distance [16], and the Aitchison distance[1], which is a Euclidean distance quantified following centered log ratio (CLR) transformation of abundances.

In addition to PAM and hierarchical clustering, we also included a third approach, the model-based Dirichlet Multinomial model (DMM) [10]. This algorithm assumes that observations of the same cluster come from the same multinomial distribution, and that parameters of the multinomial distribution have a Dirichlet distribution. This model better captures overdispersion than simply assuming a multinomial.

In this paper, we systematically compared methods for clustering microbiome observations from four published studies with either geographical or seasonal variables as the true cluster label, which enables biological interpretation of the group separation. We first applied clustering with five methods, and assessed performance of the various methods using the adjusted Rand index with the true clustering assignment. The adjusted Rand index is based on a pairwise membership agreement and is corrected for the expected value, where a score of 0 is expected with random clustering, and a score of 1 indicates perfect clustering. We then explored relationships between differences in performance with properties of the datasets considered. With the findings obtained from the method comparisons, we proposed a novel combined metric that provided high performance for all four datasets. We further test our findings with a clinical dataset where clusters are less separated.

## Results

We used four published stool microbiome datasets to test the performance of the clustering methods. A brief summary of these datasets is shown in Table 1. Each study comprises two groups of samples with relatively distinct microbiome profiles. The Shannon diversity of the dataset is calculated as the sum of Shannon diversities over all samples in the dataset, and details related to this alpha diversity are provided in a later section. The percent of sequences in high abundance OTUs is the percent of sequences in OTUs with average abundance greater than 0.001. While the individual studies varied in their 16S rRNA gene deep sequencing methods, we processed the downloaded raw sequences using the same pipeline, including VSEARCH[20] to dereplicate and remove singletons and the UNOISE2 function in usearch [6] to identify OTUs. More information about the four example datasets is provided in Supplemental Table S1. A heatmap of the high abundance OTUs can also be found in Supplemental Figure S2.

**Table 1:**
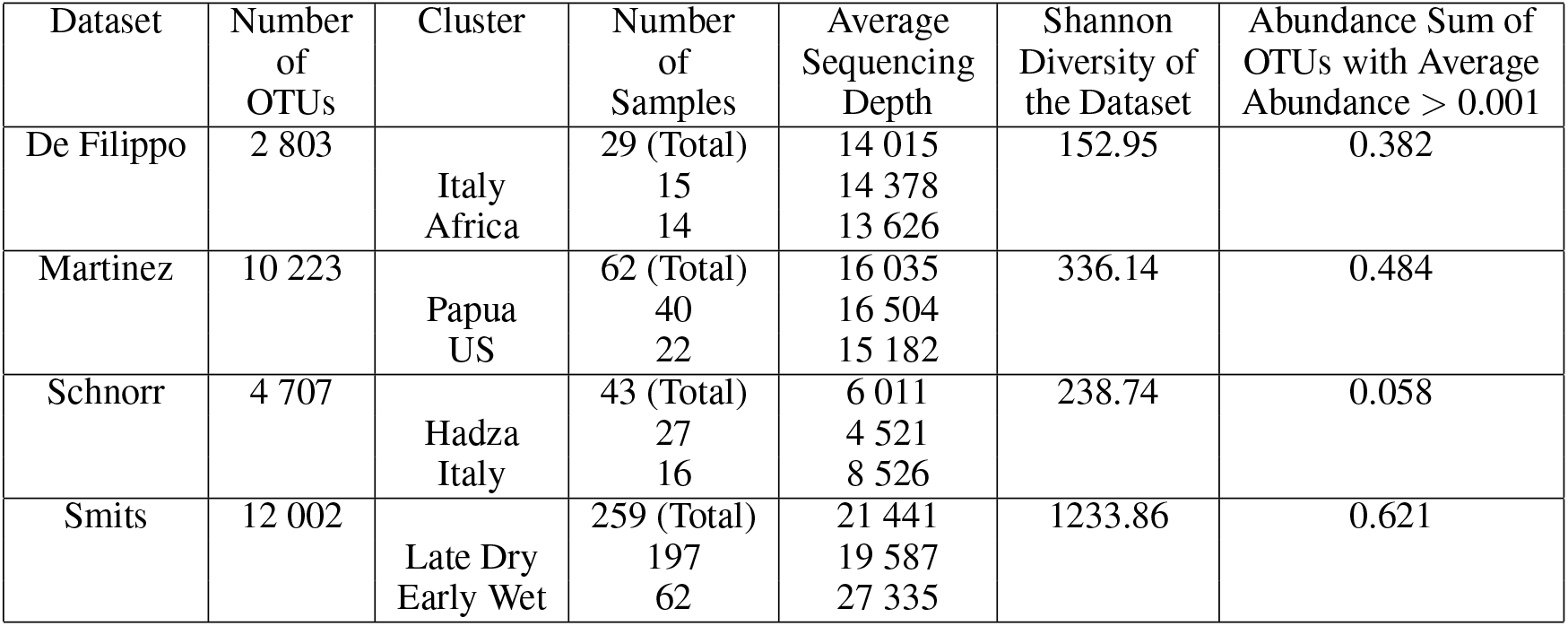
Summary of the example datasets

The first three studies compare cohorts separated by geography. The De Filippo [5] study consists of samples from healthy children living in a rural West African village in Burkina Faso, who consume a high-fiber, largely vegetarian diet, as well as samples from healthy European children of a similar age consuming a Mediterranean diet. The Martínez, et al. [17] dataset consists of samples from the Asaro and Sausi communities in Papua New Guinea, as well as those from the United States. The dataset from Schnorr, et al. [21] compares fecal samples from Italian urban adults with samples from adult hunter-gatherers living in Hadza land in Tanzania, Africa, who belong to one of the few populations that have maintained a traditional lifestyle with limited exposure to processed foods.

The fourth study, Smits et al. [25], comprises fecal samples longitudinally collected from a cohort of Hadza. Their diets are significantly affected by seasonal food availability, with more berries and honey during the wet season and more meat in the dry season. In this study, the authors found that some taxa disappear when certain foods become scarce and reappear when the seasons change. For the purpose of unsupervised learning, we used samples from the two most distinct seasons, the late dry and the early wet seasons.

The first column in Figure 1 illustrates the adjusted Rand indices of the four datasets, with different colors indicating the beta diversity metrics using PAM clustering or Dirichlet multinomial mixture model (DMM). Because of its limited capacity to handle high-dimensional data, when fitting the DMM model, we binned the OTUs present in less than 20% of the observations into one OTU when calculating Rand indices. Also, due to the fact that Dirichlet multinomial mixture is model-based, there is no corresponding distance matrix that can be shown in a PCoA plot. The black bars in the Rand index plots and the last column in the PCoA plots are generated from our newly proposed method introduced later in this paper. Due to the generally similar results between PAM and hierarchical clustering (see Figure 2), we presented results in our main text only using PAM. Among these existing methods, we found that there is no clearly superior one that universally performs well in all four datasets. Interestingly, unweighted UniFrac performs well in three of four datasets, despite lacking information regarding bacterial abundances. Similar observations for presence/absence methods have been made by Martino, et al. [18]. Moreover, the Schnorr dataset has a perfect Rand index with UniFrac distances, but a low Rand index with Bray-Curtis, Aitchison, and DMM, all of which ignore phylogenetic relationships between OTUs. We devote the next section to exploring aspects of these differences in performance. In this section, we will focus on the underperformance of Bray Curtis for the Schnorr dataset, followed by the underperformance of unweighted UniFrac for the Smits dataset. Because in reality researchers may not know the information of the true cluster assignment, we will focus on *a priori* properties of the dataset, rather than the differences between the two assigned clusters.

**Figure 1:**
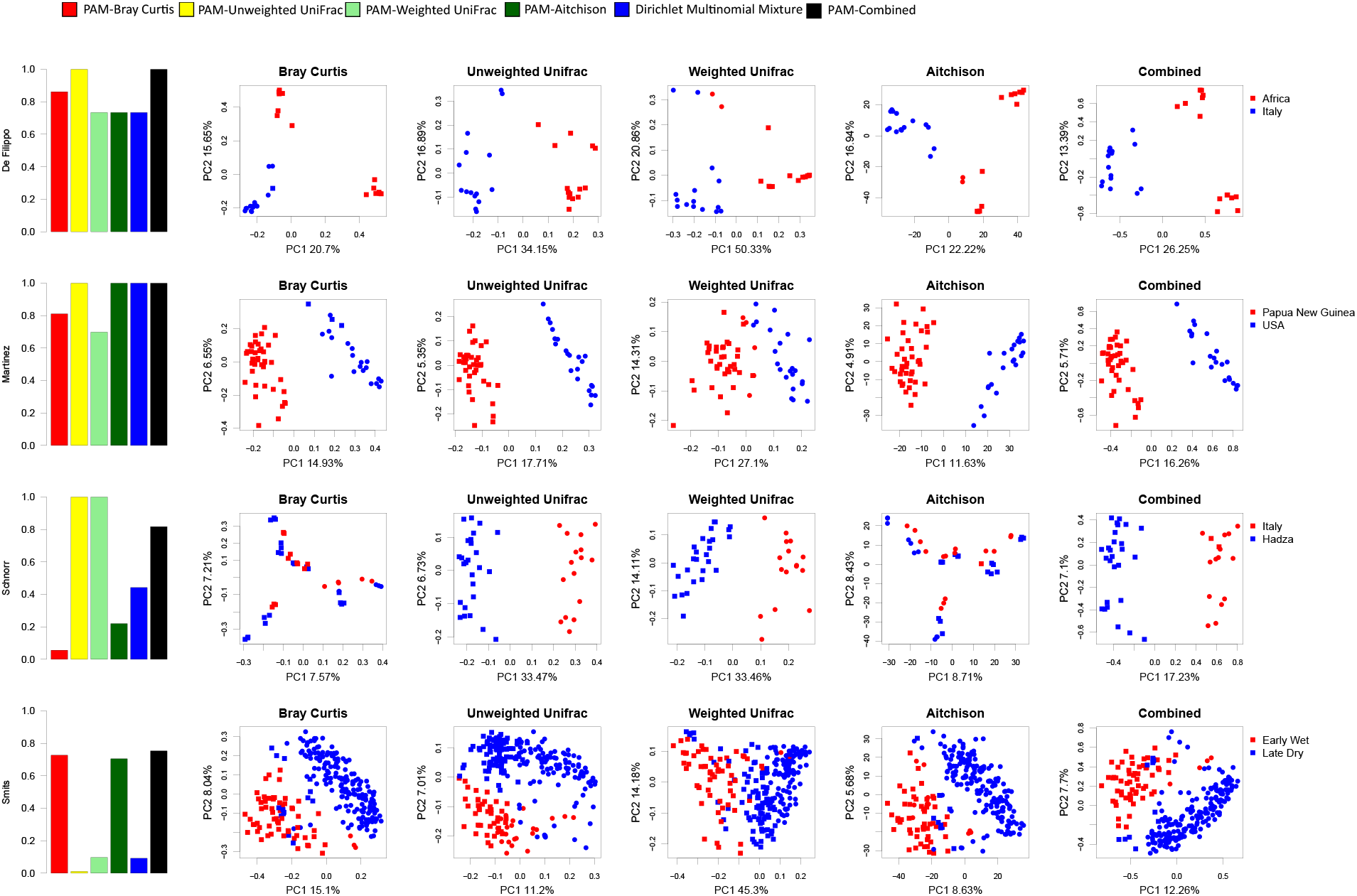
Heterogeneous Rand indices with PCoA plots of four example datasets using different methods

**Figure 2:**
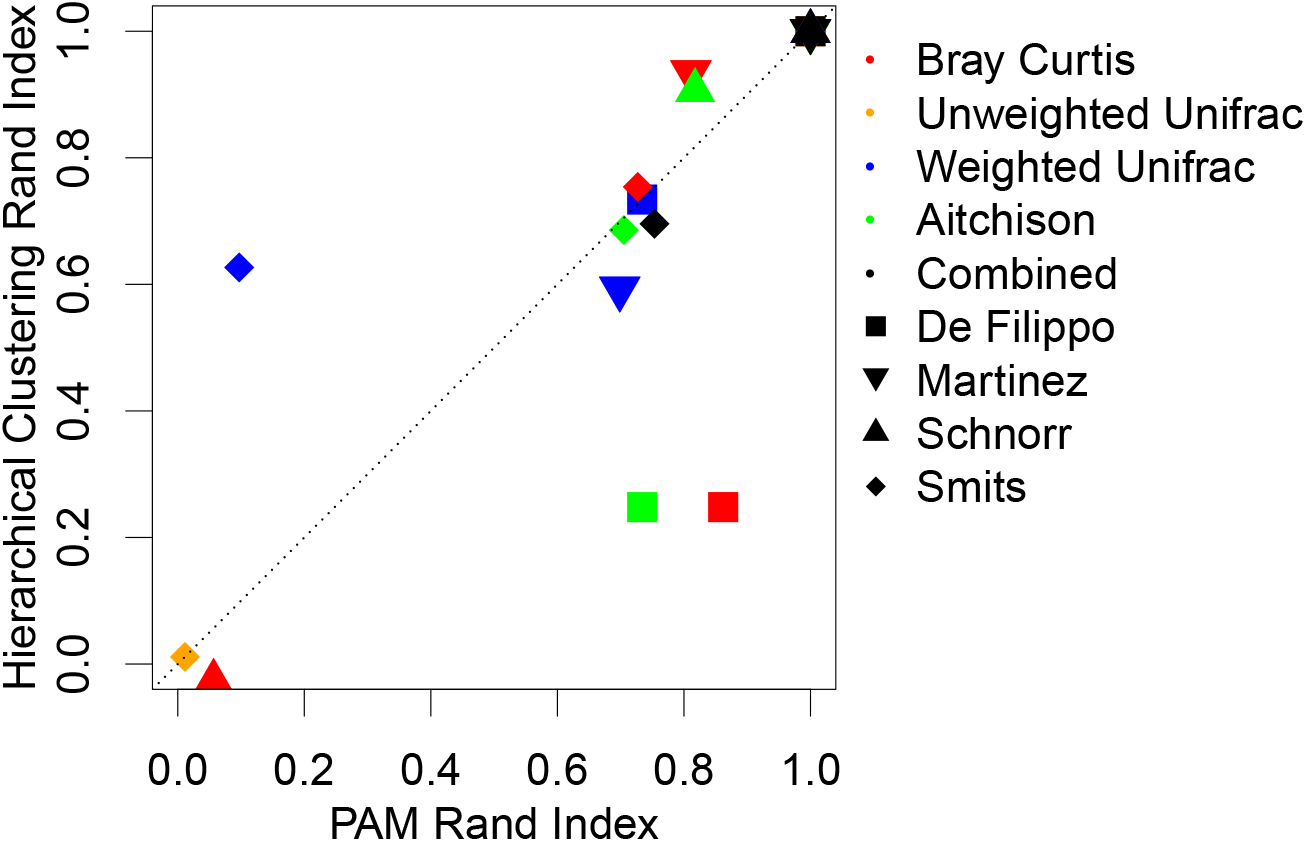
The Rand indices of PAM and hierarchical clustering correlate for the same dataset using the same beta diversity metric

### High Abundance OTUs Drive the Performance of the Bray-Curtis Dissimilarity

For a *p × n* table with *p* OTUs and *n* observations, the Bray-Curtis distance between two observations A and B is calculated with the actual counts,

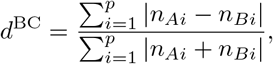

where *n*_*Ai*_ is the count for the *i*th OTU for observation *A* and *n*_*Bi*_ is the count for the *i*th OTU for observation *B*. By examining the definition, we find that the dissimilarity of the Bray-Curtis metric is driven by differences in high-abundance rather than low-abundance OTUs. Here we chose to define a metric, to potentially identify poor Bray-Curtis performance by summing the average abundance of “high abundance OTUs”, which are those with a mean abundance greater than 0.001, though similar results can be obtained across a range of thresholds.

As shown in the last column of Table 1 and the last column of the Supplemental Table S1, the sum of average abundance and the high-abundance OTUs for the Schnorr dataset is far below that of the other three datasets. A visual description is also provided as a heatmap in Supplemental Figure S2. Thus the Schnorr dataset is characterized by both unusually poor clustering performance by Bray Curtis and few high-abundance OTUs (Figure 1). To calculate this association, we attempted to improve the performance of Bray Curtis by generating novel OTUs with higher mean abundances. We did this by first generating a phylogenetic tree of the aligned OTUs, and then systematically merging OTUs starting with those most distal to the tree root. In essence, we performed “trimming” of branches from the tree, by combining the abundances of the distal OTUs to generate new, more proximal OTUs with higher mean abundances, as shown in Figure 3 (A).

**Figure 3:**
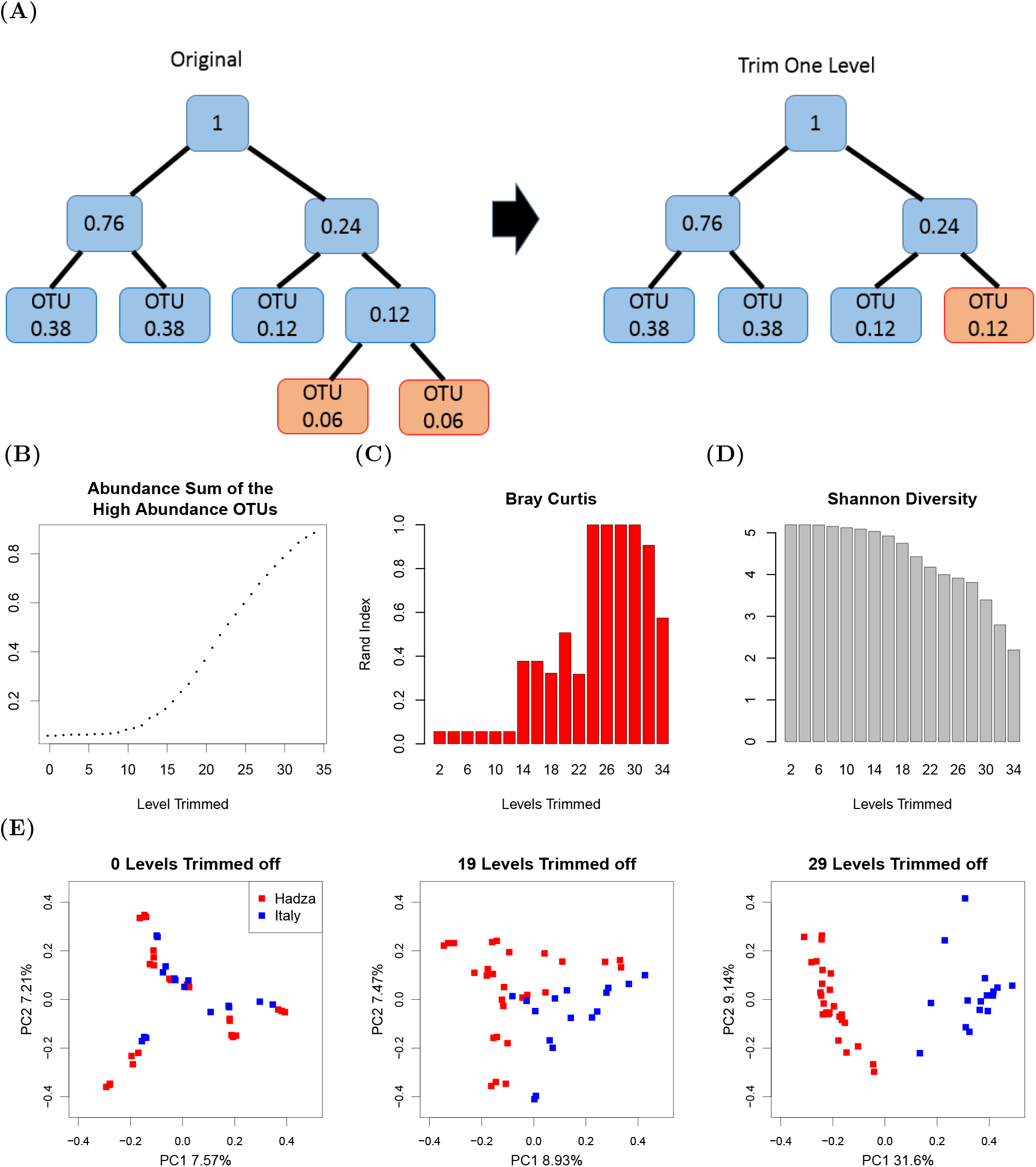
Illustration of the trimming process, summed average abundance of high-abundance OTUs plot, Rand index plot and PCoA plots of the trimmed Schnorr dataset (A) A schematic of a phylogenetic tree of OTUs, where the number in each node is the sum of the average abundance (left). After trimming the furthest branches, the average abundance of the new tree tip is greater than before (right). (B) The summed abundance of the high abundance OTUs grows with increased trimming. (C) The Rand index of Bray Curtis - PAM initially increases and then decreases with continued trimming of the phylogenetic tree towards the root. (D) The total Shannon diversity of the dataset decreases with the trimming process. (E) Bray-Curtis beta diversity PCoA plots show the separation of two natural clusters with no trimming, 19 levels of trimming, and 29 levels of trimming.

In Figures 3 (B) and (C), the X-axes indicate the number of levels trimmed, ranging from the first most distal branch to, at most, 34 levels proximal. Figure 3 (B) shows how the summed abundance of high abundance OTUs increases with more iterations of the trimming process, while Figure 3 (C) shows that the Rand index improves initially as distal sparse OTUs are merged together. As trimming continues, the Rand index begins to worsen, as presumably excessive binning results in loss of distinctive OTU information. The three PCoA plots of Figure 3 (E) correspond to the original dataset (where no levels have been trimmed), to a modified dataset where 19 levels are trimmed off, and to a modified dataset where 29 levels are trimmed off from the most distal branch towards the root. The two natural clusters begin to distinctly separate when 19 levels of branches are trimmed, and are fully separated when 29 levels are trimmed. The corresponding heatmaps of the three datasets in Supplement Figure S3 also demonstrate that trimming can increase the number of OTUs with high mean abundance.

Notably, as we progressively reduced the resolution of microbiome data during the trimming process, reflected by the decreasing Shannon diversity in Figure 3 (D), the PAM clustering showed improved performance. Shannon diversity is a commonly used alpha diversity metric used to quantify the information within a dataset. The Shannon diversity for observation *j* is defined as:

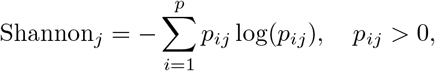

where *p*_*ij*_ is the abundance of the *i*th taxa for observation *j*. A reasonable explanation for the increase in Rand indices is that the trimming process actually utilizes the tree phylogenetic information, and similar thinking was previously mentioned in Gopalakrishnan et al. [9] and Peled et al. [19]. We encourage researchers to explore their datasets by trimming with the tools provided in our R package “MicrobiomeCluster”. Interestingly, datasets with clear clustering into two groups are likely to produce a bimodal histogram of pairwise distances between samples. One mode corresponds to small within-cluster distances, while the other mode corresponds to larger between-cluster distances. As a counterexample, we chose to examine in more detail the Martínez dataset for its high clustering performance when using the Bray-Curtis metric. We reasoned that a high degree of discriminating information was being provided by OTUs with high mean abundances. To simulate OTUs with low mean abundances progressively, we took counts assigned to an OTU and assigned those to a new OTU in randomly selected samples from each tip of the original tree to form first-generation descendants, then grew two new branches from each of the newly formed tree tips to obtain the second-generation descendants, and so on. As zero counts do not contribute to the Shannon diversity, we essentially “grew” branches without increasing the information from the original dataset, as shown in Figure 4 (A). Sequences of each OTU were randomly assigned to either of the two daughter branches. As shown in Figure 4 (B), the summed abundance of the OTUs with high mean abundance is reduced. The performance of both Bray Curtis- and Aitchison-based clustering show a decreasing trend, while tree-structure-based UniFrac methods are minimally affected in these simulations (Figure 4 (C)). With an increasing number of descendants, fewer OTUs have a high mean abundance when averaged over samples, corresponding with a drop in the performance of the Bray Curtis- and Aitchison-based methods.

**Figure 4:**
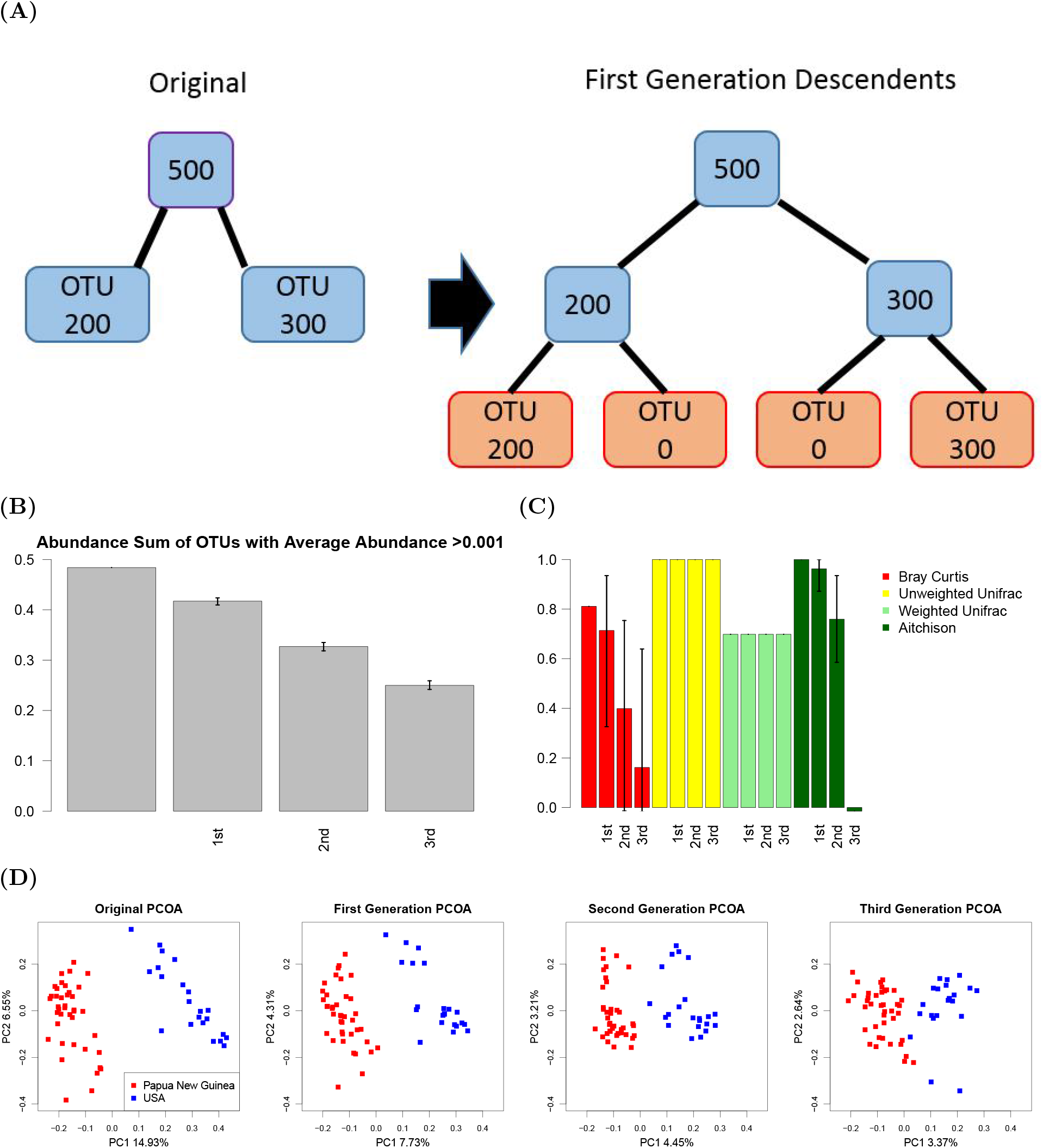
Illustration of branching process, abundance sum of OTUs with high abundance plot, Rand indexes plot, and PCoA plot of Martínez dataset with descendants (A) A schematic of the branching process, where sequences of each OTU were randomly assigned to either of the two daughter branches. (B) Abundance sum of OTUs with high mean abundance plot of the Martínez Dataset with first, second, and third generation descendants. Lines in the bar plot indicate 95th percentile intervals of the Rand indices from 200 repeated simulations. (C) Rand indices of the Martínez Dataset with the original dataset, and simulations of the first, second, and third generation descendants. Lines in the bar plot indicate 95th percentile intervals of the Rand indices from 200 repeated simulations. (D) Bray-Curtis beta diversity PCoA of the original Martínez dataset, as well as examples of datasets with the addition of the first, second, and third generation novel OTUs.

To visualize the effects of reduced OTU abundances, we plotted the PCoA of the original dataset, as well as examples of datasets with the addition of the first, the second, and the third generation simulated distal OTUs. As additional descendants are generated, the separation between two clusters becomes less apparent (Figure 4 (D)). We also plotted heatmaps of the original dataset, as well as examples of modified datasets with the addition of the first, and second generation descendants in Supplemental Figure S4. As tree branches progressively diverge, the most abundant OTUs have reduced mean abundances.

### Prevalence of Low Abundance OTUs Inhibits Clustering Performance of the Unweighted UniFrac Distance

Another key observation from our examination of clustering performance in the four datasets is that the unweighted UniFrac distance performs well for three of the datasets, but markedly underperforms in the Smits dataset. This dataset is distinct from the others in that samples were collected at different time points from the same group of individuals, rather than being collected from two distinct groups of subjects. Thus the association of the microbiome is with time (distinct in two seasons), rather than the source. We would anticipate that, during the late dry season, many bacteria more suited for the early wet season will be reduced in abundance but may not be totally eliminated and can recover when the host diet eventually becomes more hospitable, and vice versa. The geographically separated clusters from the other studies are less likely to have an extensive overlap in presence of bacterial taxa, with many taxa being present only in one population and absent in the other.

To explore determinants of when the unweighted UniFrac metric could lead to reduced clustering performance, we focused on the unweighted nature of the measure, where only the presence or absence of an OTU is quantified rather than the abundance of an OTU. In the Smits dataset, we expected that a large number of OTUs are shared by both clusters. To identify a metric that captures the information uncaptured by unweighted UniFrac, we propose using the total Shannon diversity, which is the sum of the Shannon diversities over all the samples of the dataset. Because zero entries in the OTU table do not contribute to total Shannon diversity, the total Shannon diversity in a dataset is a metric that can reflect the amount of information uncaptured by unweighted UniFrac across samples. To examine this, we simulated new repetitions of the data from the Smits dataset by gradually switching non-zero entries to 0, in essence adjusting the criteria by which an OTU would be considered “present” in a sample, as shown in Figure 5 (A). In both Figures 5 (B) and (D), the x-axis is the threshold going from 0 to 70. During the process, entries with a number of sequences less than or equal to the threshold will be converted to 0. In Figure 5 (B), the performance of unweighted Unifrac goes up with the number of entries converted to 0, whereas the performance of Bray-Curtis only fluctuates slightly. In Figure 5 (D), the total Shannon diversity drops down sharply with the number of entries converted to 0.

**Figure 5:**
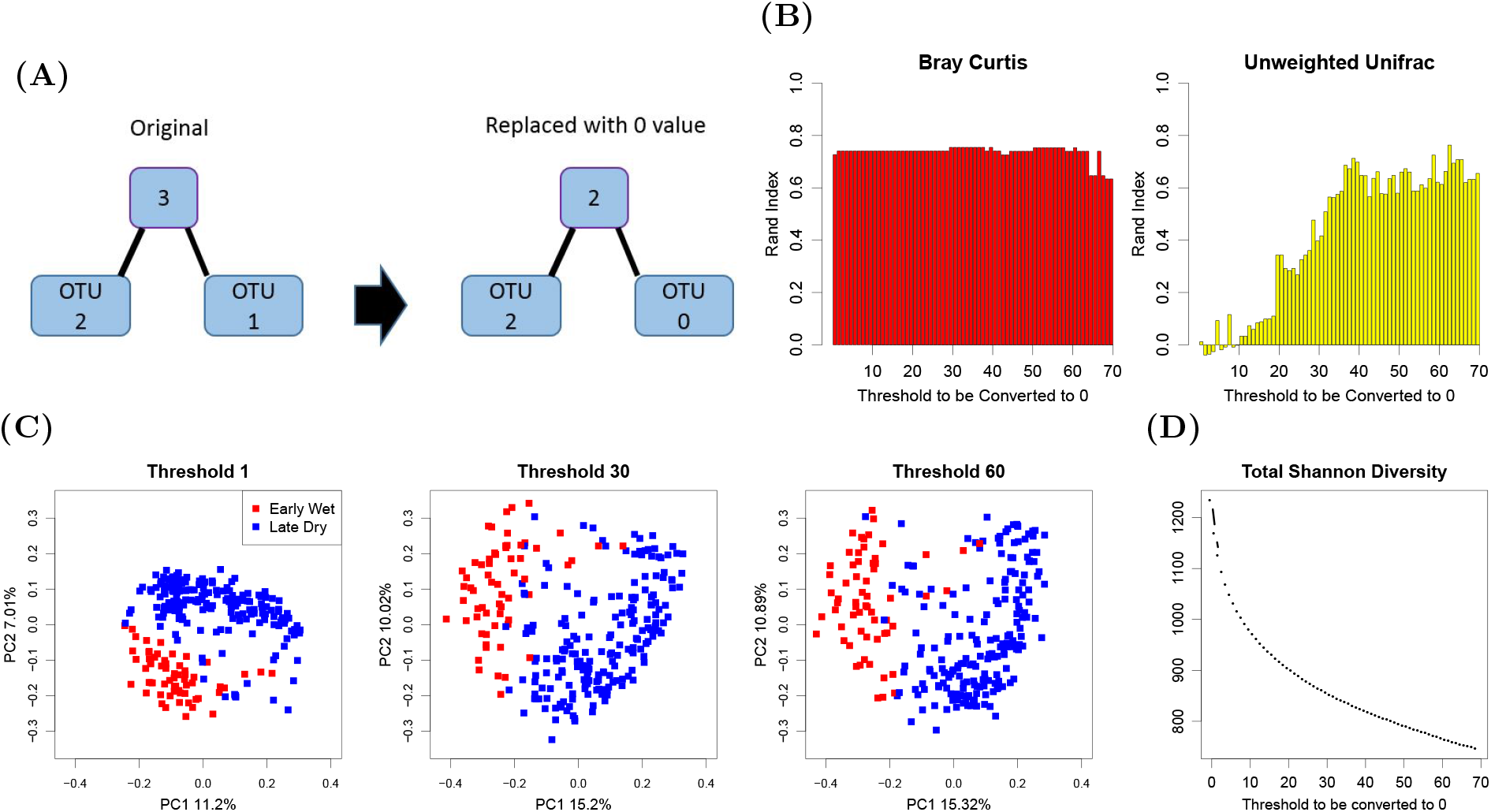
An illustration of replacing low abundance OTU with 0 value, Rand index plot, PCoA plots and the total Shannon diversity plot of the modified Smits dataset (A) A schematic plot illustrates how we replace low abundance values with a 0 value when the threshold is set to be 1. (B) Clustering performance of unweighted UniFrac improves by increasing the threshold used to change non-zero entries to 0 in the Smits dataset, while the performance of Bray Curtis remains the same. (C) Unweighted UniFrac beta diversity PCoA of the original data, the dataset where entries less than 30 are converted to 0, and the dataset where entries less than 60 are converted to 0. (D) The total Shannon diversity of versions of the Smits dataset decreases as we raise the threshold, below which counts are converted to 0.

PCoA also visually depicts improved discrimination of unweighted Unifrac with an increasing threshold. In Figure 5 (C), we plotted the PCoA of the original data, the simulated dataset where entries less than 30 are converted to 0, and the simulated dataset where entries less than 60 are converted to 0. As more non-zero entries are converted to 0, the two clusters separate further in the first coordinate. As a counter example, we chose to examine in more detail the Martínez dataset, for which the unweighted UniFrac displays high clustering performance. To simulate poorly discriminating data, we increasingly substituted 0 entries with a single count value. Such an alteration of the dataset imposes a minimal change to the original data. As with the Smits dataset, we quantified the median with the 95% empirical confidence intervals of Rand indices and Shannon diversities over 200 simulated datasets. In the x-axes of both Figure 6 (B) and (C), the probability of filling a 0-value entry with 1 goes from 0 to 1, with a 0.1 increment. Figure 6 (B) demonstrates that the total Shannon diversity indeed increases as 0-value entries are increasingly replaced with a value of 1. In turn, we found that the Rand indices of other methods are almost completely unchanged with the perturbation, while the performance of the unweighted UniFrac-based clustering is progressively reduced in Figure 6 (C).

**Figure 6:**
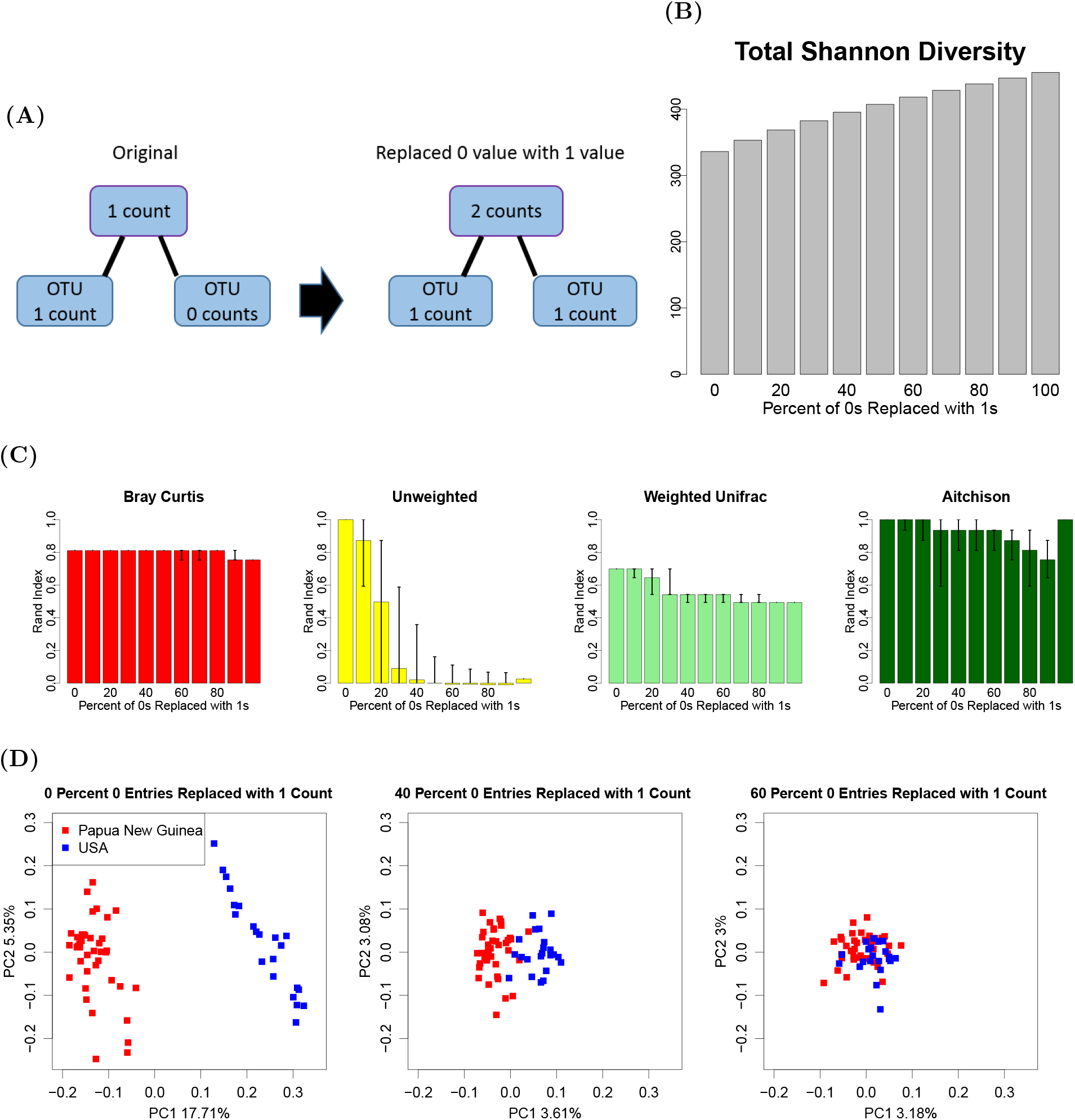
An illustration of replacing value of 0 with value of 1, the total Shannon diversity plot, Rand index plot, and the PCoA plots of the modified Martínez datasets (A) A schematic plot illustrates how we increased the low-abundance OTUs by replacing value of 0 with value of 1. (B) The total Shannon diversity increases with the number of 0 entries replaced with 1. (C) Rand index of the Martínez dataset with an increasing number of 0 entries replaced with 1. (D) Unweighted UniFrac beta diversity PCoA of Martínez dataset with 0, 40, 60 percent of 0 entries replaced with 1.

The PCoA plots of the original dataset, as well as a simulated dataset where 40 percent of 0-value entries in the OTU table are replaced with a value of 1, and a simulated dataset where 60 percent of 0-value entries are replaced with a value of 1 are shown in Figure 6 (D). We found that, as percentage of replacement increased, the two groups of samples become progressively less distinct.

### Combined Metric Shows Stable Good Performance

With the above findings, it would be appealing to develop a metric that can use the information of both high and low abundance OTUs. An intuitive way of constructing such a new metric is to combine information from both Bray Curtis and unweighted UniFrac metrics, which tend to complement each other. After normalizing pairwise Bray Curtis and unweighted UniFrac metrics by their largest values the two metrics lie between 0 and 1. We define the distance between two observations in the new metric as

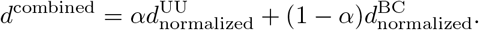

Different *α* values change the relative contributions of unweighted UniFrac and Bray Curtis distances respectively, and we suggest using *α* = 0.5 as a default simple choice. Previously, a parameter dependent metric, generalized UniFrac, was proposed by Chen et al [3]. The Supplemental Figure S5 shows that our combined metric tends to do better, and the default choice gives a robust performance across example datasets. As shown in the last column of the bar plots and the last column of PCoA plots of Figure 1, the proposed new metric (with *α* = 0.5), which inherits complementary insights from the Bray Curtis and the unweighted UniFrac dissimilarities, results in high performance in Rand indices for all four datasets.

### Evaluation in a setting without strong cluster separation

We also explored a clinical setting where one does not expect global separation between groups [9]. The study examines the relationship between microbiome and response to immunotherapy drugs for melanoma patients. Here we consider patients who responded to (30 patients) and patients who did not respond to immunotherapy drugs (13 patients) as two clusters. There are 1455 OTUs in this dataset and the average sequencing depth is 48765. Compared with four datasets discussed in previous sections, its abundance sum of OTUs with average abundance *>* 0.001 is higher (0.874), while its Shannon diversity is lower (132.75).

As shown in Figure 7, all Rand indices are less than 0.5, with the combined metric being the highest. It is interesting to notice that the combined metric can outperform each of its individual components. The Bray Curtis had the second best performance, corresponding with the presence of high abundance OTUs. Of note, clustering performance by unweighted UniFrac is poor, and the low Shannon diversity of the dataset would normally indicate the potential for better performance.

**Figure 7:**
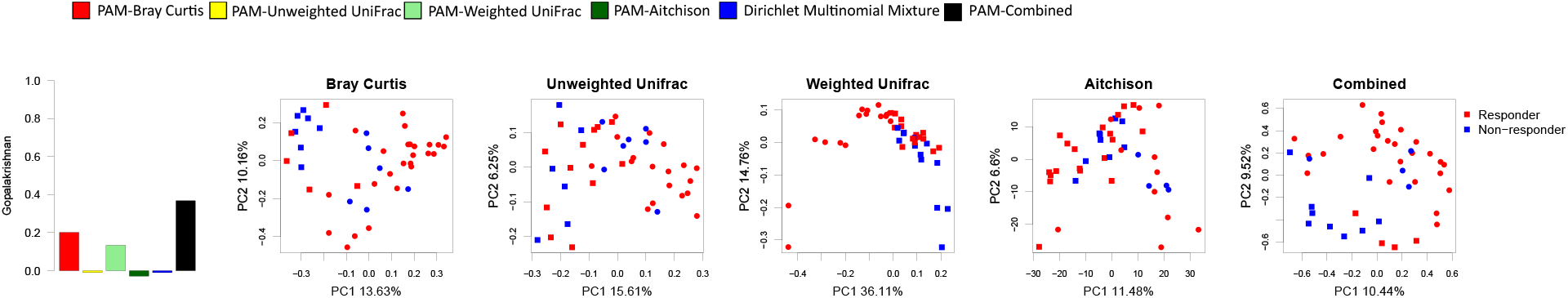
Rand indices plot and PCoA plots under various metrics for Gopalakrishnan dataset (A) The Rand indices plot. (B) The PCoA plot under 5 metrics.

## Discussion

A quantitative recommendation regarding when to avoid certain metrics would require a systematic examination of many more datasets. We did observe that in four of the datasets we considered, hierarchical clustering, though less frequently used with microbiome data, can provide similar and sometimes superior results compared to the more commonly used PAM method. (Figure 2) Unlike PAM clustering, the number of clusters do not have to be pre-determined prior to hierarchical clustering, which is a potential advantage. For supervised settings where the group membership is known, beta diversity metrics, including the proposed combined beta metric, can be used as input to distance-based testing approaches such as PERMANOVA. For the data sets considered here, PERMANOVA produces significant p-values across virtually all metrics and data sets. This is because PERMANOVA takes the group labels as known, and tests the null hypothesis that both the centroids and dispersion are the same across groups. There may be evidence to reject this null even in settings where the groups overlap substantially.

## Conclusions

Our systematic evaluation of clustering performance in these five datasets show that there is no existing clustering method that universally performs the best across all datasets. In general, the weighted UniFrac outperforms other methods across studies. When OTUs with high mean abundance are rare, clustering methods that do not consider phylogeny (i.e., Bray-Curtis, Aitchison, DMM) perform less well. In contrast, in a dataset with lots of low abundance OTUs, the unweighted UniFrac tends to produce poor separation, while other methods can still perform well. To capitalize on the complementary strength of the two metrics, we propose a new metric that incorporates the Bray Curtis metric and the unweighted UniFrac metric, which gives good performance in a dataset representative of the common scenario where clusters are less separated.

## Declarations

### Ethics approval and consent to participate

Not applicable

### Consent for publication

Not applicable

### Availability of data and material

We created an R package “MicrobiomeCluster,” which contains methods implemented in this manuscript (https://github.com/YushuShi/MicrobiomeCluster.git) [23], including the proposed combined metric as well as some basic data manipulation tools.

### Author’s contributions

RRJ designed the research. YS conducted the data analysis and wrote the manuscript, with input from LZ, CBP and KAD. RRJ, CBP and KAD revised the manuscript. All authors read and approved the final manuscript.

## Acknowledgements

Not applicable

## References

[1] J. Aitchison. Principal component analysis of compositional data. Biometrika, 70(1):57–65, 1983.

[2] J. R. Bray and J. T. Curtis. An ordination of the upland forest communities of southern Wisconsin. Ecological monographs, 27(4):325–349, 1957.

[3] J. Chen, K. Bittinger, E. S. Charlson, C. Hoffmann, J. Lewis, G. D. Wu, R. G. Collman, F. D. Bushman, and H. Li. Associating microbiome composition with environmental covariates using generalized Unifrac distances. Bioinformatics, 28(16):2106–2113, Aug. 2012.

[4] M. Claesson, I. Jeffery, S. Conde, S. Power, E. O’Connor, S. Cusack, H. M B Harris, M. Coakley, B. Lakshminarayanan, O. O’Sullivan, G. F Fitzgerald, J. Deane, M. O’Connor, N. Harnedy, K. O’Connor, D. O’Mahony, D. Van Sinderen, M. Wallace, L. Brennan, and P. W O’Toole. Gut microbiota composition correlates with diet and health in the elderly. Nature, 488:178–84, 07 2012.

[5] C. De Filippo, D. Cavalieri, M. Di Paola, M. Ramazzotti, J. B. Poullet, S. Massart, S. Collini, G. Pieraccini, and Lionetti. Impact of diet in shaping gut microbiota revealed by a comparative study in children from Europe and rural Africa. Proc Natl Acad Sci U S A, 107(33):14691–14696, 2010.

[6] R. C. Edgar. UNOISE2: improved error-correction for Illumina 16S and its amplicon sequencing. bioRxiv, 2016.

[7] J. Fukuyama. Emphasis on the deep or shallow parts of the tree provides a new characterization of phylogenetic distances. Genome Biology, 20(131), 2019.

[8] J. A. Gilbert, M. J. Blaser, J. G. Caporaso, J. K. Jansson, S. V. Lynch, and R. Knight. Current understanding of the human microbiome. Nat Med, 24(4):392–400, 2018.

[9] V. Gopalakrishnan, C. Spencer, L. Nezi, A. Reuben, M. Andrews, T. Karpinets, P. Prieto, D. Vicente, K. Hoffman, S. Wei, et al. Gut microbiome modulates response to anti-pd-1 immunotherapy in melanoma patients. Science, 359(6371):97–103, 2018.

[10] I. Holmes, K. Harris, and C. Quince. Dirichlet multinomial mixtures: Generative models for microbial metagenomics. PLOS ONE, 7(2):1–15, 02 2012.

[11] J. Jovel, J. Patterson, W. Wang, N. Hotte, S. O’Keefe, T. Mitchel, T. Perry, D. Kao, A. L. Mason, K. L. Madsen, et al. Characterization of the gut microbiome using 16S or shotgun metagenomics. Frontiers in microbiology, 7:459, 2016.

[12] L. Kaufman and P. J. Rousseeuw. Partitioning around medoids (program pam). Finding groups in data: an introduction to cluster analysis, pages 68–125, 1990.

[13] R. Knight, C. Callewaert, C. Marotz, E. R. Hyde, J. W. Debelius, D. McDonald, and M. L. Sogin. The microbiome and human biology. Annual review of genomics and human genetics, 18:65–86, 2017.

[14] O. Koren, D. Knights, A. Gonzalez, L. Waldron, N. Segata, R. Knight, C. Huttenhower, and R. E. Ley. A guide to enterotypes across the human body: Meta-analysis of microbial community structures in human microbiome datasets. PLOS Computational Biology, 9(1):1–16, 01 2013.

[15] C. Lozupone and R. Knight. Unifrac: a new phylogenetic method for comparing microbial communities. Appl Environ Microbiol, 71(12):8228–8235, 2005.

[16] C. A. Lozupone, M. Hamady, S. T. Kelley, and R. Knight. Quantitative and qualitative β diversity measures lead to different insights into factors that structure microbial communities. Appl Environ Microbiol, 73(5):1576–1585, 2007.

[17] I. Martínez, J. C. Stegen, M. X. Maldonado-Gómez, A. M. Eren, P. M. Siba, A. R. Greenhill, and J. Walter. The gut microbiota of rural Papua New Guineans: composition, diversity patterns, and ecological processes. Cell reports, 11(4):527–538, 2015.

[18] C. Martino, J. T. Morton, C. A. Marotz, L. R. Thompson, A. Tripathi, R. Knight, and K. Zengler. A novel sparse compositional technique reveals microbial perturbations. mSystems, 4(1), 2019.

[19] J. U. Peled, S. M. Devlin, A. Staffas, M. A. Lumish, R. Khanin, E. R. Littmann, L. Ling, S. Kosuri, M. A. Maloy, J. Slingerland, K. F. Ahr, K. A. P. Rodriguez, Y. Shono, A. E. Slingerland, M. Docampo, A. D. Sung, D. Weber, A. M. Alousi, B. Gyurkocza, D. M. C. Ponce, J. Barker, M.-A. Perales, S. A. Giralt, Y. Taur, E. G. Pamer, R. R. Jenq, and M. R. M. van den Brink. Intestinal microbiota and relapse after hematopoietic-cell transplantation. Journal of Clinical Oncology, 35 15:1650–1659, 2017.

[20] T. Rognes, T. Flouri, B. Nichols, C. Quince, and F. Mahé. VSEARCH: a versatile open source tool for metagenomics. PeerJ, 4:e2584, 2016.

[21] S. L. Schnorr, M. Candela, S. Rampelli, M. Centanni, C. Consolandi, G. Basaglia, S. Turroni, E. Biagi, C. Peano, M. Severgnini, et al. Gut microbiome of the Hadza hunter-gatherers. Nature communications, 5:3654, 2014.

[22] C. E. Shannon. A mathematical theory of communication. Bell System Technical Journal, 27(4):623–656, 1948.

[23] Y. Shi. MicrobiomeCluster, 2020. R package.

[24] E. H. Simpson. Measurement of diversity. Nature, 163(688), April 1949.

[25] S. A. Smits, J. Leach, E. D. Sonnenburg, C. G. Gonzalez, J. S. Lichtman, G. Reid, R. Knight, A. Manjurano, J. Changalucha, J. E. Elias, M. G. Dominguez-Bello, and J. L. Sonnenburg. Seasonal cycling in the gut microbiome of the Hadza hunter-gatherers of Tanzania. Science, 357(6353):802–806, 2017.

