## Supplemental Material for "Performance Determinants of Unsupervised Clustering Methods for Microbiome Data"

April 8, 2021

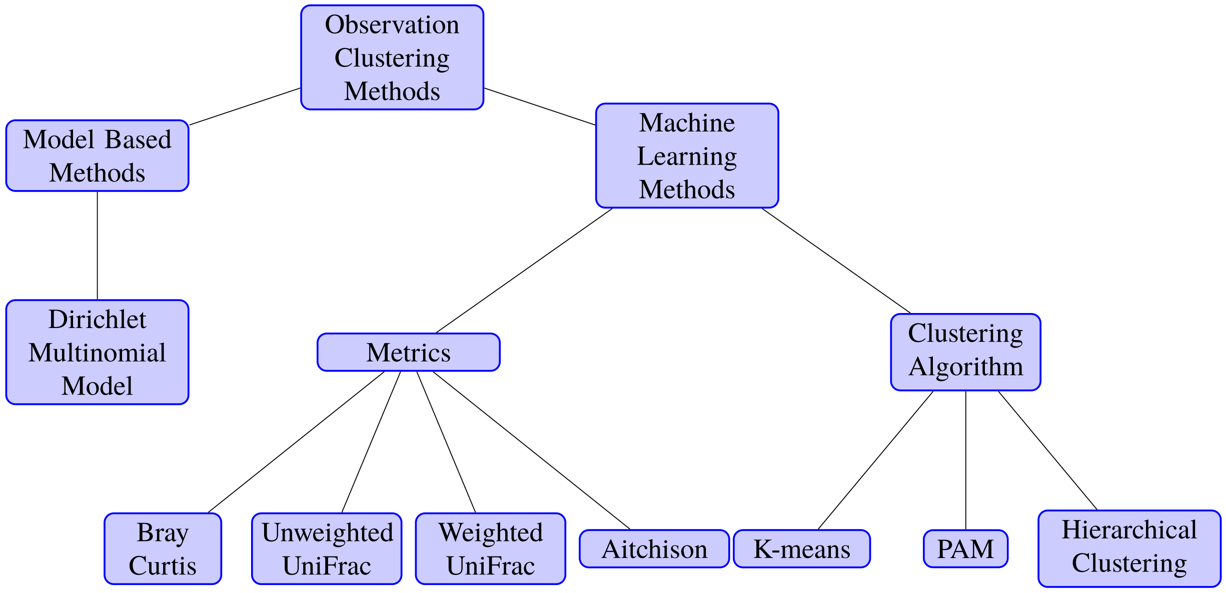


Figure S1: An illustrative plot of commonly used clustering methods

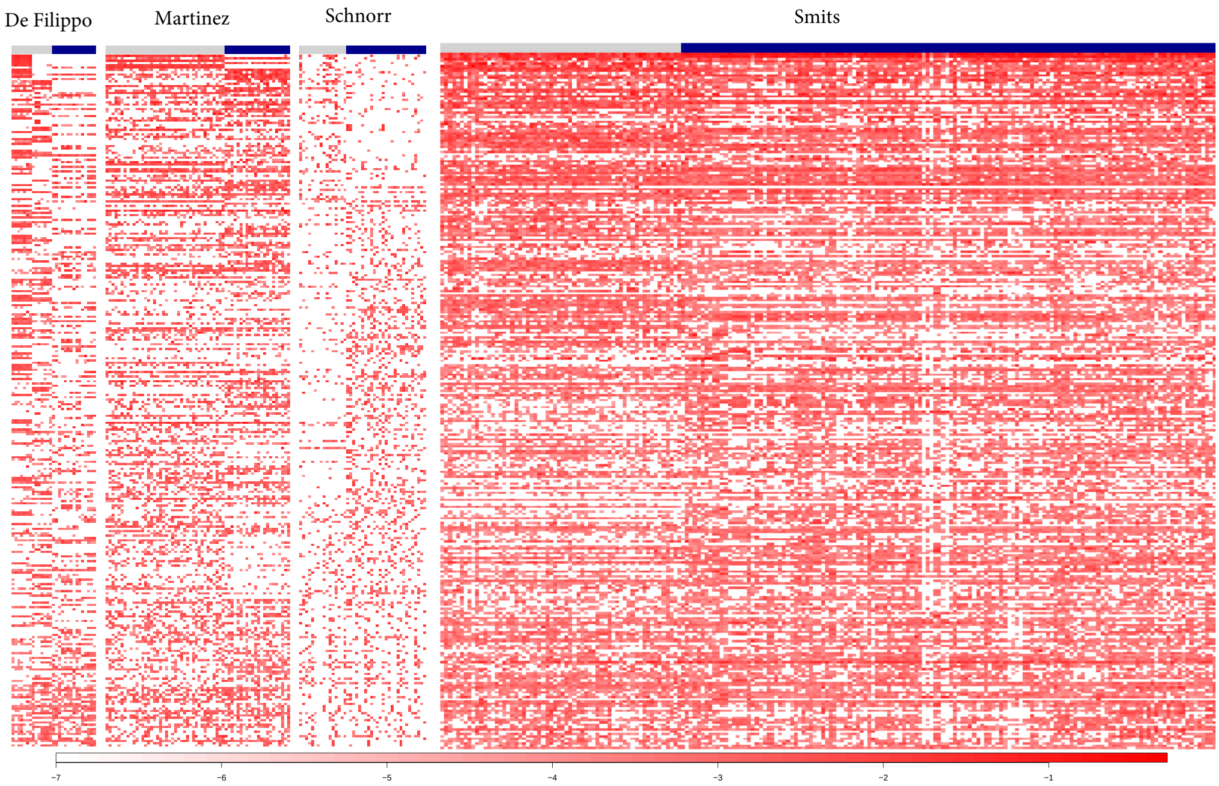


Figure S2: Heatmap of the most abundant 300 OTUs for the four example datasets
This figure shows the most abundant 300 OTUs for the four published datasets plotted in log 10 scale. Red color indicates high abundance, whereas white color indicates low abundance. As shown in the plot, the abundances of “high abundance" OTUs in Schnorr dataset are lower than those of the other three datasets. Gray and blue indicate different clusters in each dataset.

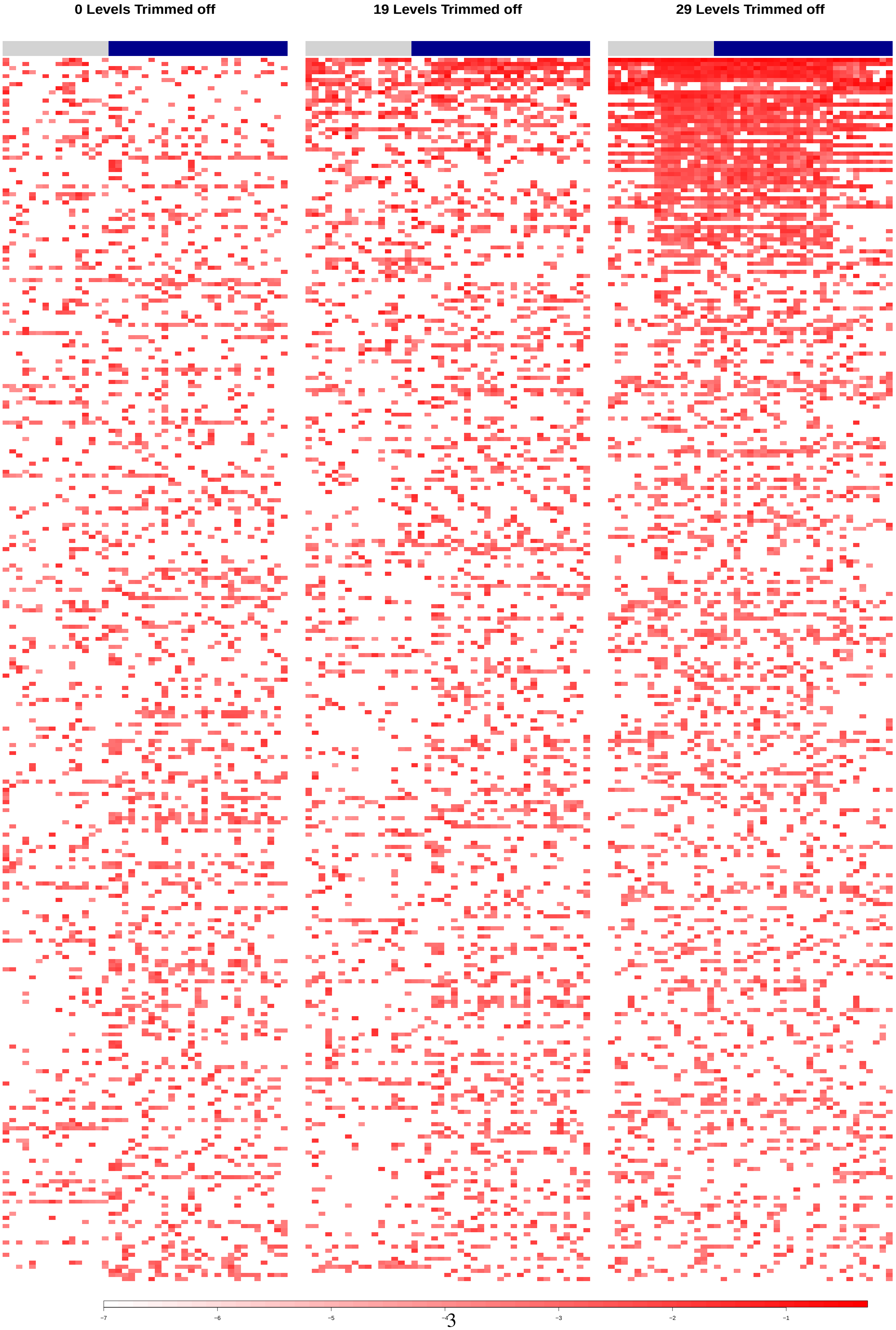




Figure S3: Heatmap of the most abundant 300 OTUs for the Schnorr dataset with 0, 19, and 29 levels trimmed off

This figure plots the most abundant 300 OTUs of the Schnorr Dataset with 0, 19, and 29 levels trimmed off. As trimming goes along, the abundant OTUs aggregate sequences from distant OTUs. In each subpanel, the gray left part is the Italian sample set, while the blue right part is the Hadza sample set.

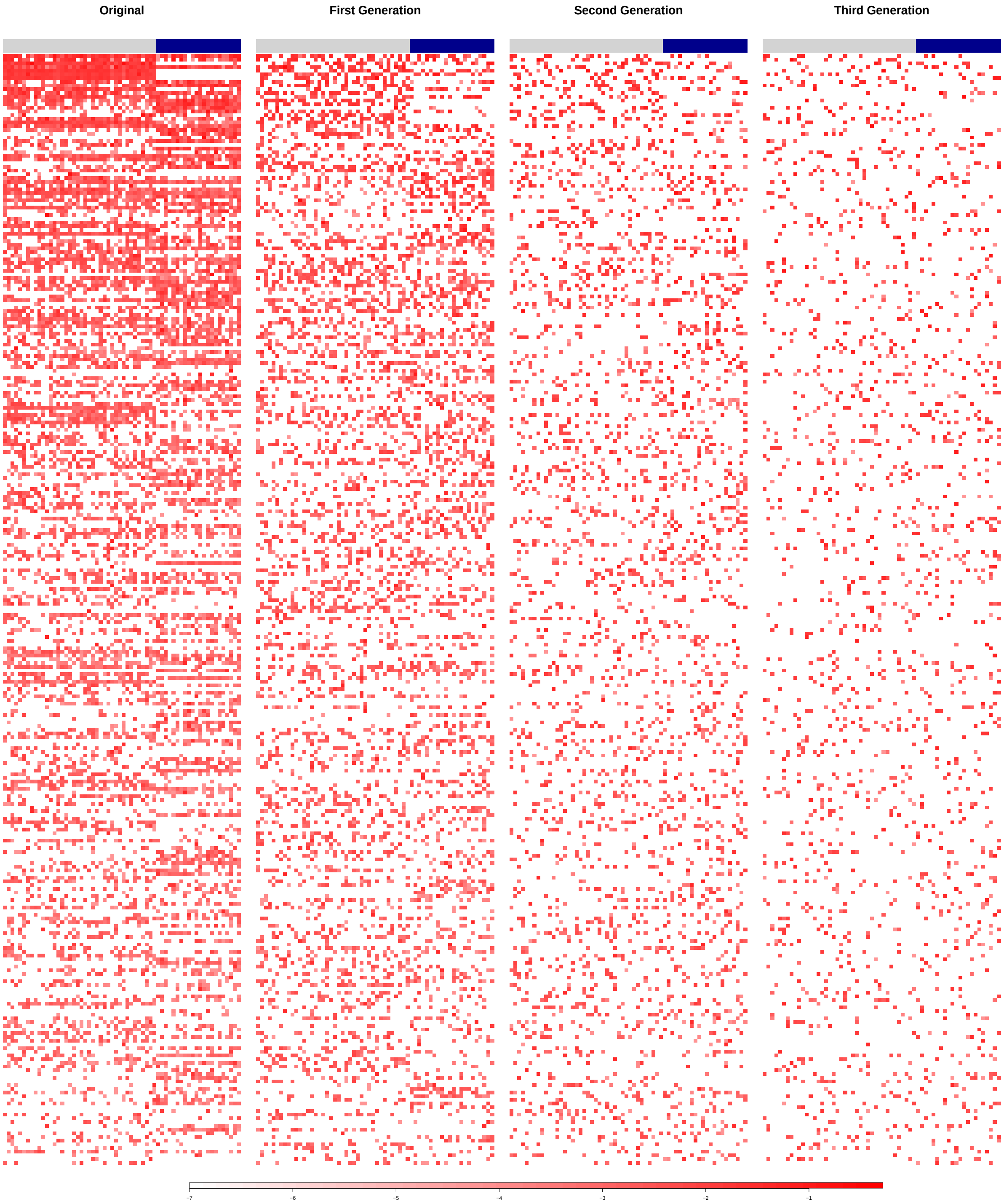


Figure S4: Heatmap of the most abundance 300 OTUs of the Martínez dataset, with descendants

This figure plots the most abundance 300 OTUs of the Martínez Dataset with its first, second and third generation descendants. As the tree branches diverge, fewer sequences are left in the most abundant 300 OTUs. In each subpanel, the gray left part is the Papua sample set, while the blue right part is the US sample set.

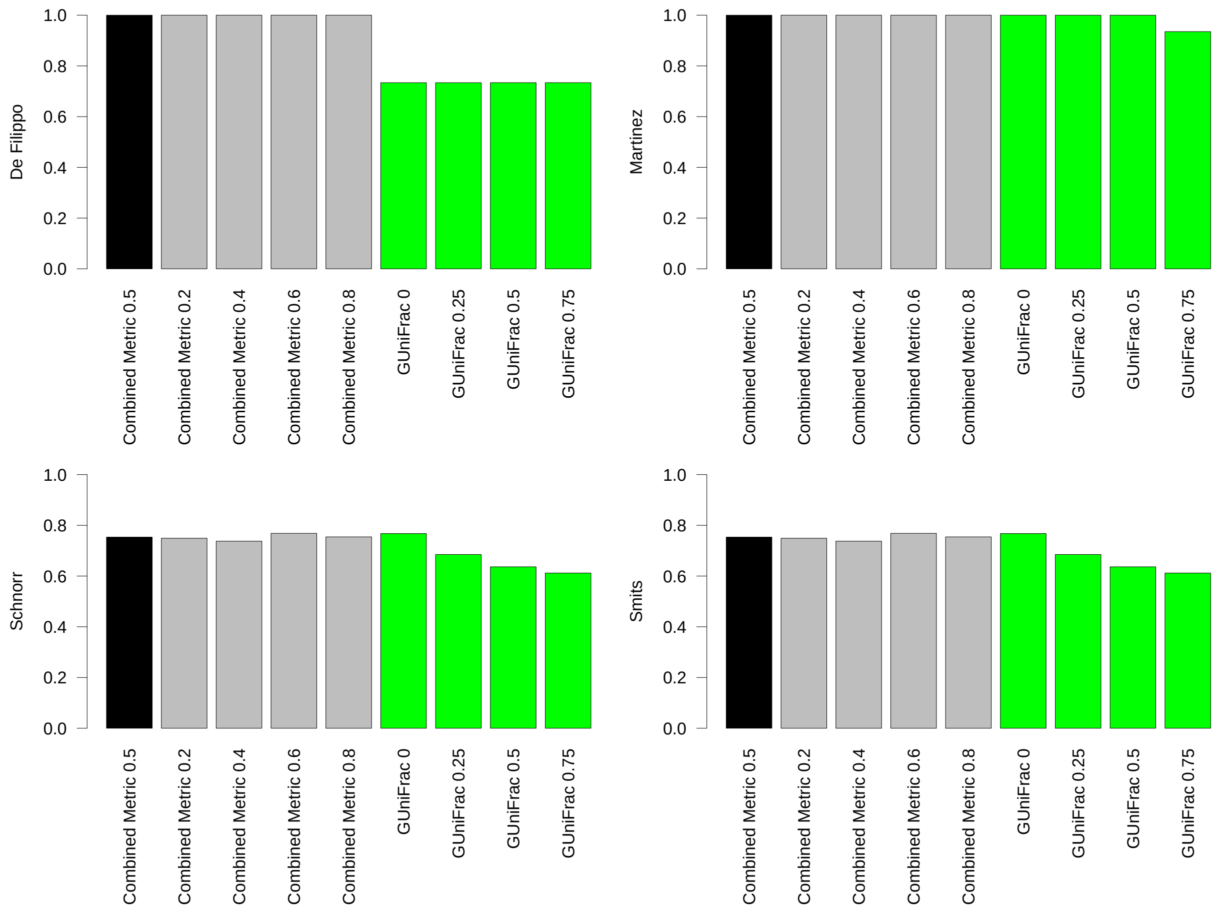


Figure S5: Rand indices with different $\alpha$ values for the proposed metric and comparison with the generalized UniFrac metric

This figure shows the Rand indices with different values of $\alpha$ (0.2, 0.4, 0.6, 0.8) for the proposed metric and compares the results with the Generalized UniFrac under different parameters for it (0, 0.25, 0.5, 0.75).

The third and fourth columns of this table refer to the skewness and kurtosis of the averaged abundance vector of the whole dataset or each cluster. Skewness is often used as a measure of symmetry, while kurtosis is a measure for the degree of tailedness in the distribution. Prevalence of high abundance OTUs is the percent present in samples for OTUs with an averaged abundance greater than $0.001$.

| Dataset | Cluster | Skewness | Kurtosis | Percent of 0 Entries | Prevalence of High Abundance OTUs |
|---|---|---|---|---|---|
| De Filippo | | 6.39 | 72.19 | 77.2 | 33.3 |
| | Italy | 5.61 | 49.38 | | |
| | Africa | 8.77 | 128.53 | | |
| Martínez | | 23.91 | 780.56 | 83.2 | 53.46 |
| | Papua | 27.44 | 961.11 | | |
| | US | 20.45 | 513.49 | | |
| Schnorr | | 3.49 | 27.13 | 88.2 | 19.14 |
| | Hadza | 2.52 | 15.12 | | |
| | Italy | 5.50 | 62.75 | | |
| Smits | | 63.23 | 4649.79 | 85.7 | 76.15 |
| | Late Dry | 65.11 | 4805.62 | | |
| | Early Wet | 37.40 | 1778.04 | | |

Table 1: Additional Summary of the Example Datasets
